# Deep Mutational Scanning Reveals a De Novo Disulfide Bond and Combinatorial Mutations for Engineering Thermostable Myoglobin

**DOI:** 10.1101/2024.02.24.581358

**Authors:** Christoph Küng, Olena Protsenko, Rosario Vanella, Michael A. Nash

## Abstract

Engineering protein stability is a critical challenge in biotechnology. Here, we used massively parallel deep mutational scanning (DMS) to comprehensively explore the mutational stability landscape of human myoglobin (hMb) and identify key mutations that enhance stability. Our DMS approach involved screening over 10,000 hMb variants by yeast surface display, single-cell sorting and high-throughput DNA sequencing. We show how surface display levels serve as a proxy for thermostability of soluble hMb variants, and report strong correlations between DMS-derived display levels and top-performing machine learning stability prediction algorithms. This approach led to the discovery of a variant with a *de novo* disulfide bond between residues R32C and C111, which increased thermostability by >12 °C compared to wild-type hMb. By combining single stabilizing mutations with R32C, we engineered combinatorial variants that exhibited predominantly additive effects on stability with minimal epistasis. The most stable combinatorial variant exhibited a denaturation temperature exceeding 89 °C, representing a >17 °C improvement over wild-type hMb. Our findings demonstrate the capabilities in DMS-assisted combinatorial protein engineering to guide the discovery of thermostable variants, and highlight the potential of massively parallel mutational analysis for the development of proteins for industrial and biomedical applications.

## Introduction

In protein engineering, understanding the intricate relationships between sequence, structure and function is paramount for identifying variants with enhanced properties. A common approach uses laboratory directed evolution ^1–4^, however, success rates are low and the discovery process is stochastic. Especially in campaigns for improving enzyme activity, limits are imposed by the natural trade-offs between stability and phenotypic activity ^5^. This universal phenomenon often leads to the destabilization of proteins when introducing mutations that improve function. A strategy to overcome or attenuate this effect is to start with a highly stable variant that can tolerate destabilizing but functionally beneficial mutations.

It was shown, for example, through lattice simulations and experiments that highly stable sequences are more amenable to *in vitro* evolution and can achieve higher levels of functional improvement upon mutation^6^. This suggests a generalizable strategy for protein engineering involving the design of superior variant libraries that are biased to preferentially include stabilizing mutations^7^. In the past, approaches based on random mutagenesis^8^ or ancestral sequence reconstruction^9^ have been used for improving the thermostability of enzymes. A more recent approach used large-scale mutational data to create mutability landscapes and identify regional trends in mutational tolerance^10,11^. Such an approach allows to systematically screen mutational neighborhoods to find hotspots of stabilizing mutations, as well as sequence stretches where mutations are detrimental to stability. In this context, massively parallelized methods for assaying thousands of variants in parallel as is typically done in mutational scanning (DMS) experiments can unlock valuable information for informed library design^12,13^.

In this work we use a DMS workflow to study human myoglobin (hMb), a 17 kDa globular heme protein that serves as an ideal model for our study highlighting structural features important for stability. In addition to its native function as an oxygen binding protein, hMb has been investigated as an engineering scaffold for the introduction of new catalytic activity. For instance, Hilvert and colleagues used directed evolution to improve the peroxidase activity of myoglobin^14^. Other campaigns transformed hMb into an artificial metalloenzyme capable of performing atom transfer radical cyclisation as well as keto reductase catalysis^15,16^.

Here, we quantified expression levels of yeast surface displayed holo hMb variants by DMS. We show how expression levels can serve as a proxy for thermostability and reveal mutational effects on hMb stability in a massively parallel workflow. We note that in literature there is extensive evidence that yeast surface display expression level is generally correlated with protein folding stability^5,17–19^. Thermodynamically destabilizing mutations yield lower detectable expression levels in yeast surface display because less stably folded domains are susceptible to yeast quality control mechanisms which hinder the secretion of unstable variants. Although mRNA stability, protease susceptibility and other factors may also influence surface display expression levels, we consider the estimated fitness scores as relevant to thermodynamic folding stability, and further validated this experimentally.

We found strong correlations between our DMS dataset and hMb evolutionary conservation scores, as well as stability prediction algorithms. This approach enabled discovery of highly stabilizing mutations in hMb, including a *de novo* disulfide bonded variant. By combining single mutants, we were able to further enhance stability, observing primarily additive effects with low epistasis. This method paves the way for more efficient protein engineering of hMb catalytic activity by providing a stable scaffold for further optimization.

## Results

### DMS data generation and validation

To generate the DMS dataset, we used nicking strand mutagenesis and constructed a pooled gene library comprising 10,467 hMb variants that were expressed and displayed as Aga2p fusion proteins on the surface of EBY100 yeasts^20^. Our library comprised 2,578 single amino acid substitutions (80% of all possible), 2,769 double mutations (< 0.1% of all possible) and small fractions of higher order mutants. We measured the expression level of each variant by quantitative high-throughput fluorescence activated cell sorting (FACS) combined with NGS (**Figure 1)**. Yeast cells displaying hMb variants were stained with a fluorescent antibody in proportion to hMb expression level. The pooled library was then sorted into four bins (**Figure 1B**) and the gene sequences of variants in each bin were determined by NGS. The frequencies of variant reads in the bins were transformed into expression fitness scores using a custom computational pipeline employing tools from^21^ and ^22^ (see methods, **Tables S1-5**).

**Figure 1.**
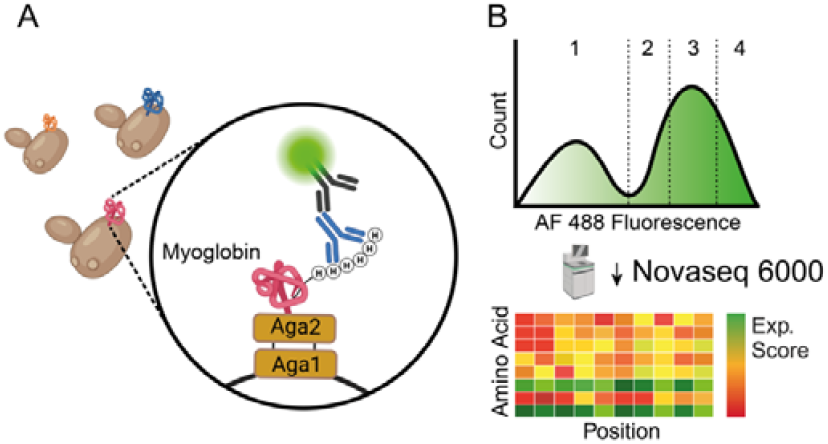
Scheme of the experimental and computational workflow. **A)** A myoglobin variant library is displayed on the surface of yeast cells and fluorescently labeled based on expression level of the hMb variant via C-terminal His_6_-Tag. **B)** The labeled library is sorted into four bins with known median fluorescence intensities. After sequencing all variants in each bin by NGS, we convert read counts into expression fitness scores to obtain the fitness landscape.

To assess reproducibility of experimental DMS fitness scores, we performed two independent biological replicates and evaluated their correlation (**Figure 2A**). These data showed strong agreement between replicates (*r* = 0.90, Pearson’s correlation coefficient p-value < 0.0001) with higher confidence for variants that were covered by a larger number of sorted cells (**Figure 2A**, color scale). As further validation, we analyzed the distribution of expression fitness values of nonsense (stop codon) and synonymous mutations (**Figure 2B**), and found that nonsense mutations (**Figure 2B**, red) exhibited low expression fitness scores (−0.52 ± 0.13, n = 126). Synonymous gene variants (**Figure 2B**, blue) that encoded WT hMb using alternate codons exhibited expression fitness values of 0.03 ± 0.07 (n = 109) on average, indicating the relevant range of fitness values for synonymous codon substitutions.

**Figure 2.**
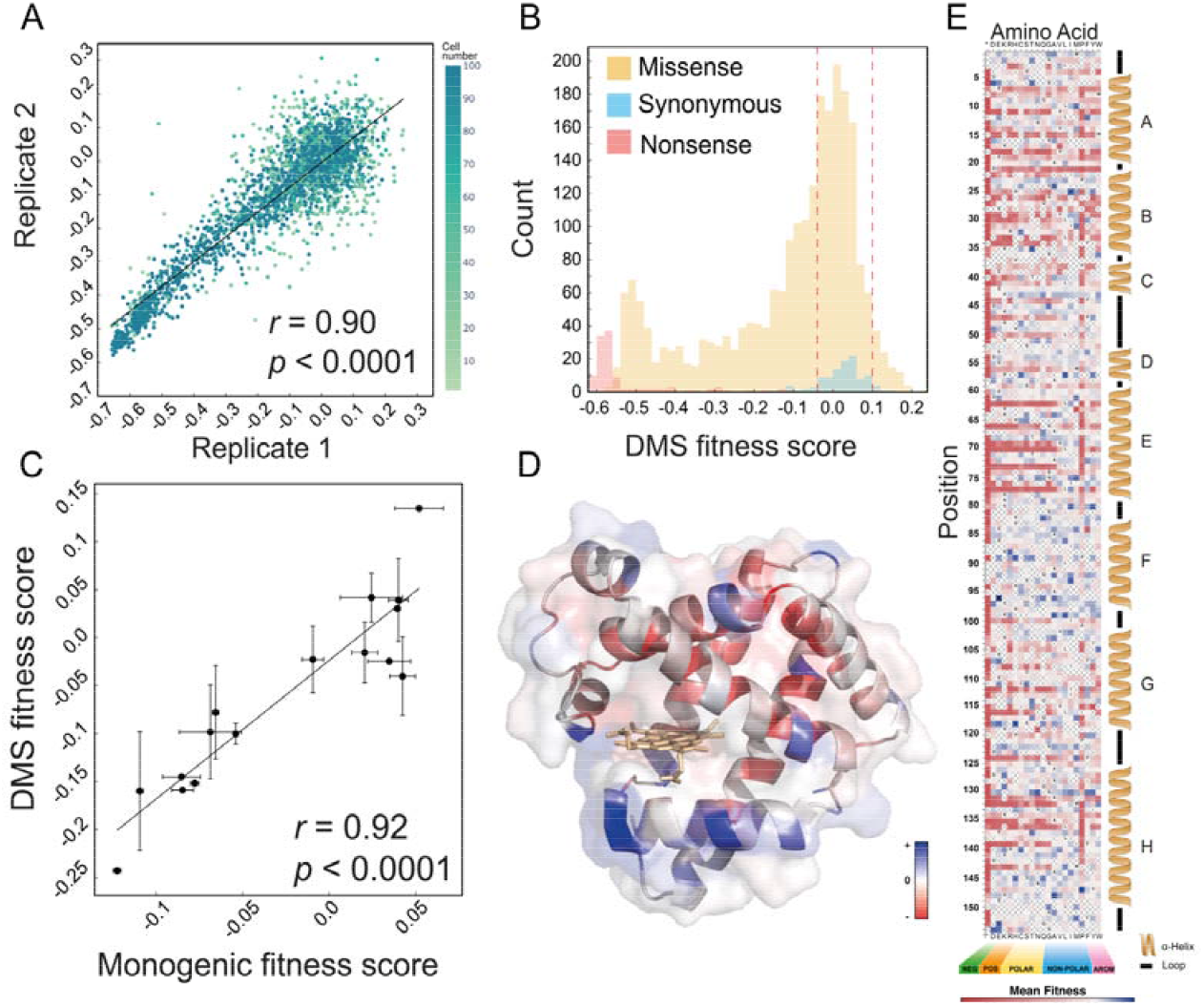
DMS of yeast displayed human myoglobin. **A)** Linear correlation of fitness scores from two biological replicates. The color gradient represents the number of cells carrying a particular hMb variant found in both replicates. **B)** Distribution of fitness scores for nonsense (red), missense (orange) and synonymous single mutations (blue). Red dashed lines indicate the width of the fitness range for synonymous mutations. **C)** Correlation of expression fitness scores determined by DMS with monogenetic scores, experimentally determined for 16 single variants. Y error bars correspond to standard deviation (Std) of grouped amino identities, X error bars correspond to Std of biological replicates. **D)** hMb 3D structure (PDB: 3RGK) colored by mean fitness score per position. **E)** Heatmap showing expression fitness scores of single substitution mutants in the dataset. Amino acids along the X-axis are ordered by biochemical properties. Black letters within the plot indicate WT residue identity at the respective position. X symbols indicate missing variants or variants with too few reads to pass the quality threshold.

Missense mutations encoding single substitutions generated a bimodal expression fitness histogram (**Figure 2B**, orange) across a broad range of values (fitness range: - 0.60 to 0.26). Most single missense mutations (67.15%) exhibited negative fitness values, indicating expression levels lower than WT hMb. To further validate expression fitness scores derived from DMS, we selected 16 variants covering a range of positive and negative fitness values in the DMS dataset. We independently synthesized their genes and tested expression fitness (F_exp_) in monogenic cultures. The observed fluorescence intensity distribution following antibody staining was then used to calculate a single variant expression score (**Figure 2C**) using the same procedures as for the *en masse* DMS experiments. We found that monogenic expression scores were strongly correlated with those measured by pooled DMS (*r* = 0.92, *p* <0.0001). Taken together, these controls demonstrate that fitness scores determined by DMS report with high fidelity on the expression levels of thousands of hMb variants in parallel.

We next visualized the per position average fitness score on the 3D structure of hMb (**Figure 2D**) and the single mutant fitness values in the form of a heatmap (**Figure 2E**). In the heatmap we noticed conspicuous dark red horizontal stripes with semi-regular periodicity within specific regions (residues 7-41, 62-77, 104-120 and 128-147), indicating that residues intolerant for mutations were found with regular spacings. After aligning these patterns with secondary structure features or 3D structure, we found that these corresponded to residues along the sides of alpha helices that faced inwards towards the hydrophobic core of the protein. By fitting a sinusoidal function to the experimental fitness scores for residues 70-79 (Helix E), we obtained a periodicity of 3.59 residues, which matched the periodicity of typical protein α-helices and further confirmed the origins of the periodicity of low fitness values (**Figure S1**). Another observation that emerged from the heatmap was the intolerance for proline mutations within helices, which manifested as a vertical dark stripe covering large ranges of residues. In the heatmap (**Figure 2E**), the amino acid substitutions are ordered by biochemical properties along the x-axis. This depiction further helps to illustrate an intolerance for polar side chains at residues facing the core of the protein. The higher-than-average tolerance of helix F to mutations is indicative of ligand (heme) binding site. Helix F is not part of the hydrophobic core, but was shown to exhibit collective motion driving ligand entry and exit^23^. In the expression score it is tolerant for mutations indicating a stability-activity trade off for hMb binding oxygen *via* the heme group.

### Myoglobin is more tolerant of hydrophobic mutations

We analyzed the effects of each endpoint amino acid (regardless of position) on fitness scores for all missense mutations. The average fitness of all missense mutants was -0.13, and we scored endpoint amino acid fitness as a percent change from this global average. This allowed assessing the influence of side chain biochemical properties on expression fitness (**Figure S2**). Apart from stop codons, the strongest negative impact was observed for mutations to proline (Avg. score: -163%, n = 122), a well known helix-breaking substitution^24^. Charged amino acids (D, E, K, R and H) were found to severely negatively influence fitness scores (Avg. score: -38%, n = 580). The myoglobin tertiary structure is mostly defined by α-helices built around a strongly hydrophobic core, and we found the least disruptive mutations to be hydrophobic residues (A, V, L, I and M), (Avg. score: +63%, n = 639). The aromatic hydrophobic side chain of phenylalanine also had higher than average fitness (Avg. score: +42%, n = 116). Many of the observed effects have been discussed in a meta-study worth mentioning analyzing the roles of amino acids among different proteins^25,26^.

### Discovery of R32C disulfide bonded myoglobin with enhanced thermostability

We found that cysteine substitutions were less detrimental to expression on average (Avg. score: +30%, n = 133). One cysteine mutation in particular (R32C) emerged with a fitness score among the highest in the entire dataset (R32C fitness = 0.13). Mapping the position of this mutation onto the 3D structure of myoglobin revealed that the original arginine side chain situated in helix B faced a native cysteine (Cys111) in the adjacent helix G. We hypothesized that the R32C mutation had introduced a new disulfide bond that increased protein stability and thereby expression level. To investigate this further, we expressed and purified R32C myoglobin from an oxidizing *E. coli* strain (Origami 2). Comparative SDS-PAGE analysis of WT and R32C myoglobin under reducing and non-reducing conditions supported the presence of the disulfide bond (**Figure 3A**). To further evaluate the presence of this disulfide bond, we analyzed purified samples of WT and R32C variants with mass spectrometry, comparing the R32C variant again under reducing and non-reducing conditions (**Figure 3C**). The increase in detected mass of 2 Daltons for the reduced sample corresponding to the two hydrogen atoms further confirmed the formation of the cysteine-cysteine bond. We used nano differential scanning fluorimetry (NanoDSF) to measure the denaturation temperature (T_m_), and found a significant increase of +12.5 °C for the disulfide variant (R32C T_m_= 83.8 °C; WT T_m_= 71.3 °C, **Figure S3**). Disulfide bond engineering as well as cross-linking of non-canonical amino acids have both been reported for modulating myoglobin properties^27,28^, however, to our knowledge the R32C myoglobin variant discovered by our DMS screen has not been described in literature.

**Figure 3.**
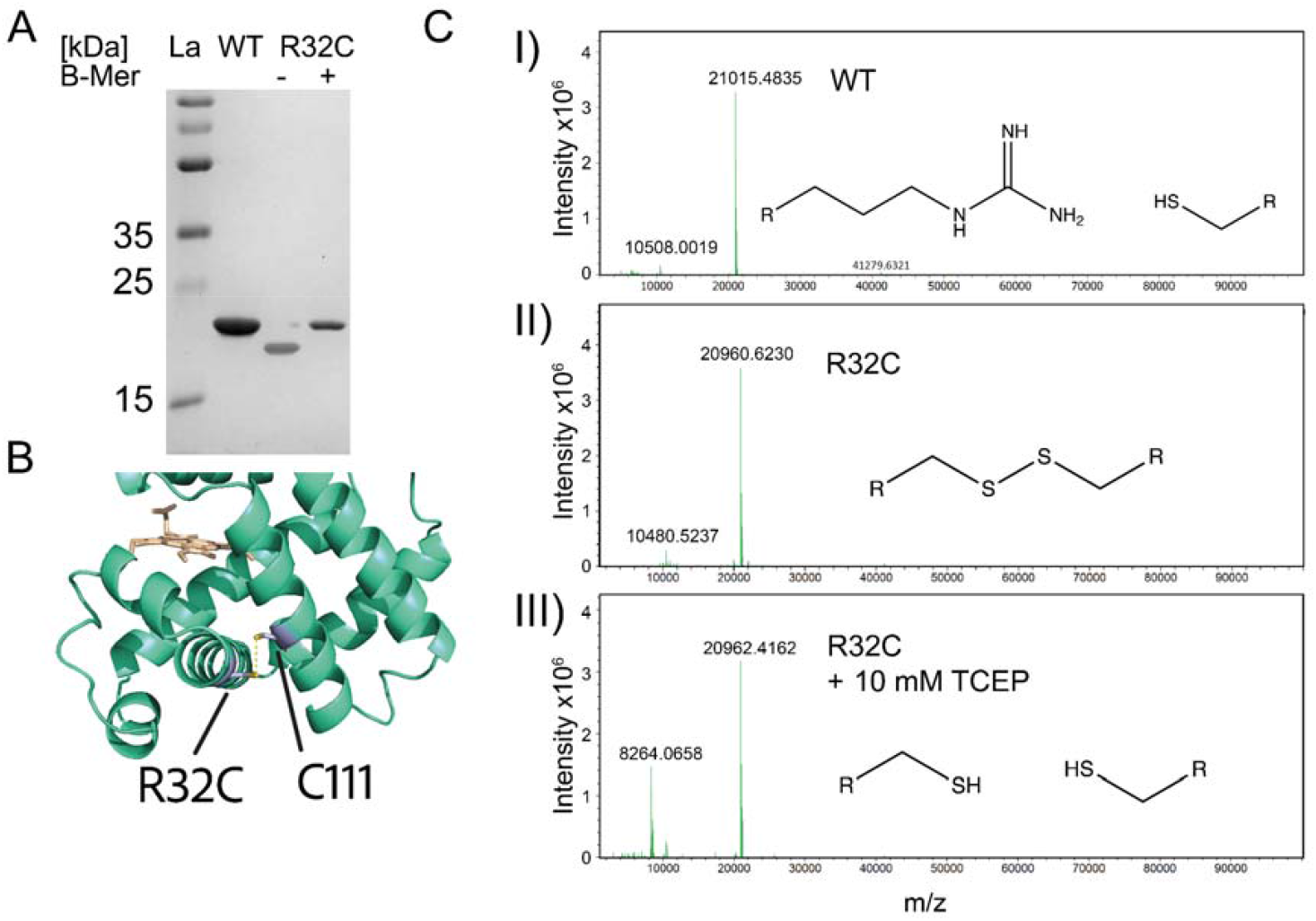
**A)** 15% SDS Page gels of purified proteins for WT and R32C. Lane 1) reference ladder; lane 2) WT hMb; lanes 3 & 4) R32C hMb variant boiled without (−) or with (+) 5% (vol./vol.) beta-mercaptoethanol reducing agent, respectively. The lower apparent molecular weight for the non-reduced sample is consistent with an intramolecular disulfide bond. **B)** Residues involved in disulfide bond formation, as mapped on protein structure (PDB: 3RGK) Cysteines are colored blue and thiols are shown in yellow. Heme cofactor is shown in wheat color. **C)** Mass spectrometry results. Deconvoluted peaks for purified WT **(I)** and R32C **(II)** protein variants. **III)** Same sample as B), incubated with 10 mM TCEP prior to desalting and MS analysis. Reduction of disulfide bond yielded a mass increase of ∼2 Daltons.

### Sequence conservation and solvent accessible surface area correlate with expression fitness

We next analyzed correlations between sequence conservation and per position average expression fitness scores. We used ConSurf^29^ to identify homologous sequences and determine conserved positions in myoglobin. The ConSurf tool used HMMER^30^ to collect 1,425 myoglobin homologues from the UNIPROT database, of which 1,360 sequences passed similarity and coverage thresholds. Out of those, 300 sequences with the lowest E-value (matches with highest significance hence excluding paralogues) were used to determine per position conservation scores on a scale from 1 to 9. We plotted the per position average fitness of all mutants in the DMS dataset vs. per position sequence conservation and found a significant negative correlation (**Figure S4 A**, (*r* = -0.92, p <0.0001), indicating more conserved positions were less tolerant for mutation on average and resulted in lower expression levels. It is important to stress that sequence conservation in myoglobin is driven not only by expression stability, but also by its function as an oxygen reservoir in skeletal and cardiac muscle tissue. Positions 46, 98, 99 and 154 surrounding the heme binding site are crucial for myoglobin function and highly conserved (CONS_Score_ = 9), however, they are highly tolerant for mutations in our expression screen, exhibiting neutral fitness or positive fitness in the case of position 99. These sites contributed to substantial variability of the negative trend (**Figure S4 A**). These data highlight the ability of DMS to shed light onto activity-stability trade-offs^5^.

Many conserved residues were found in the hydrophobic core of hMb, which prompted us to investigate correlations between expression fitness scores and solvent accessible surface area (SASA). SASA scores were extracted using Pymol from an AlphaFold model of human myoglobin (AF-P02144-F1)^31^. We assigned every residue a score between 0 and 100, corresponding to the minimum and maximum solvent exposure per position in myoglobin, respectively. The analysis revealed a positive correlation (*r* = 0.61, *p* <0.0001, **Figure S4 B**) with a prominent outlier being Asp21 (SASA_Score_ = 47), which joins helix A and B (**Figure S4 C**). We found this position does not accept any other amino acid, including negatively charged glutamate as can well be observed in **Figure 2E**. In addition, we were interested in how well yeast display expression level measured by DMS could be correlated with thermodynamic stability of soluble versions of the same protein sequences. We predicted the mean per position ΔΔG_folding_ of myoglobin (PDB: 3RGK, n = 149) using the FoldX stability prediction algorithm^32^. Comparing the predicted values to the DMS expression fitness scores, we found strong negative correlation when averaged per position (**Figure 4A**, *r* = -0.56, *p* < 0.0001) but also for single individual mutations (**Figure 4B**, *r* = -0.43, *p* < 0.0001), again validating the experimental setup serving as a proxy for thermodynamic stability. We further performed the same analysis with a more recently developed stability prediction tool (ThermoMPNN)^33^. This deep neural network builds on the ProteinMPNN framework ^34^ and was trained on a mega-scale dataset^35^ to predict change of stability in point mutations. It has been benchmarked against state-of-the-art methods and showed strong performance, especially when predicting stabilizing mutations. Again, we tested averaged and single mutation correlations with predicted ΔΔG_folding_ from ThermoMPNN and obtain Pearson correlation scores even higher than with FoldX (**Figure 4C**, *r* = -0.74, *p* < 0.0001) for averaged as well as for singles (**Figure 4D**, *r* = -0.71, *p* < 0.0001).

**Figure 4.**
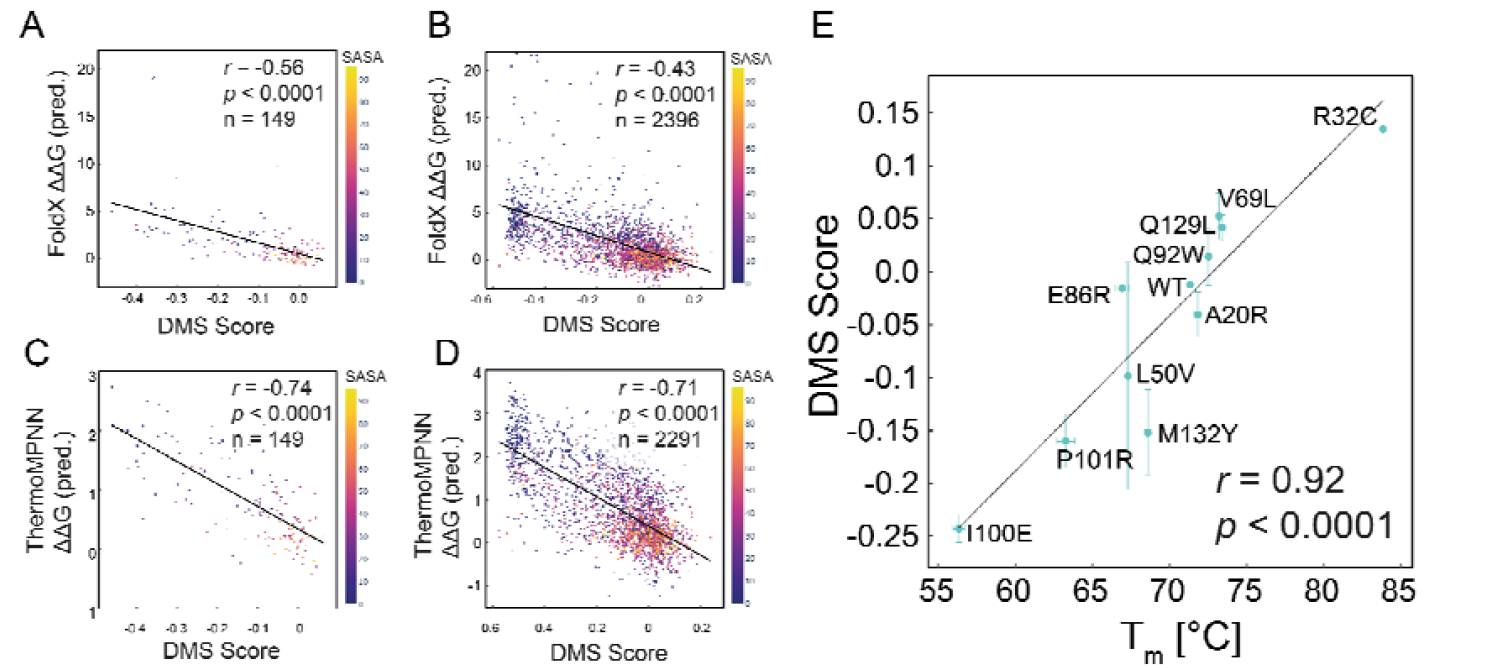
DMS scores correlate with computationally predicted and experimentally measured thermodynamic stability. **A)** Scatterplot of FoldX ΔΔG_folding_ vs. experimental DMS scores, grouped by amino acid residue position. Color coding represents solvent accessible surface area (SASA). **B)** FoldX ΔΔG_folding_ vs. DMS scores for individual variants. **C)** ThermoMPNN ΔΔG_folding_ vs. DMS scores, grouped by amino acid residue position. **D)** ThermoMPNN ΔΔG_folding_ vs DMS scores for individual variants. E) Linear correlation of DMS fitness score and melting temperature T_m_ for 10 single variants along with wild type.

To further evaluate this connection, we expressed and purified 10 of the single mutant variants used for monogenetic DMS score validation along with the WT sequence as soluble proteins in *E. coli*, and assayed thermostability by nano differential scanning fluorimetry (NanoDSF). We found strong correlation between T_m_ values measured by NanoDSF and DMS expression fitness scores (**Figure 4C**, *r* = 0.92, *p* < 0.0001).

Together these data demonstrate that on average across the mutational ensemble, the DMS expression scores from yeast surface display are predictive of thermodynamic folding stability for hMb. The strong connection between yeast expression level and protein stability has been reported in prior work and is further verified here by correlating DMS expression fitness scores with the thermostability of soluble hMb variants^19,36^. Similarly, it has been shown that in *E. coli*, expression levels of holo myoglobin highly depend on apomyoglobin stability^37^.

### DMS scores in hMb show low levels of epistasis

Next we investigated double (n = 2,769) and triple (n = 391) mutants present in the library. We processed them alongside the single mutants in the computational pipeline and filtered them for the same quality thresholds. We concluded that since most mutations are detrimental for expression level, the fraction of double mutants with lower expression than WT level would be even higher. In fact, we found that 83.90 % of the double mutants and 90.03 % of the triple mutant variants were expressed at lower levels than wild type (fitness score < 0) (Figure 5A). When comparing the DMS fitness scores of double (F_1,2_) and triple (F_1,2,3_) mutants with the sum fitness scores of the individual single mutants (i.e. F_1_ + F_2_; F_1_ + F_2_ + F_3_), we found that they correlated strongly ((**Figure 5B**, *r* = 0.87, *p* < 0.0001) for double mutants and (**Figure 5C**, *r* = 0.80, *p* < 0.0001) for triple mutants). From that observation, we concluded that in a simple protein like hMb, for low level combinations of up to three mutations, we do not observe significant epistasis. For the analysis for plots in **Figure 5**, we filtered the data for a minimal sum of F_1,2_ and triple F_1,2,3_ of -0.62, because our computational pathway, depending on the WT fitness allows for DMS fitness scores only to reach within -0.62 to 0.38. Due to the nature of the log2 operation on min-max normalized values, the data is limited to that range. While not a problem when analyzing single mutants or comparing scores, for the described analysis of linear additivity we need to take this into account, as computational scores F_1,2_ and F_1,2,3_ plateau at -0.62 while the sum of their individual mutations F_1_,F_2_ and F_3_ can result in lower scores. We note that the number of variants in **Figure 5B** and **C** differ from **Figure 5A** because we could only analyze multiple mutants for which we have the corresponding single mutants passing the quality threshold.

**Figure 5.**
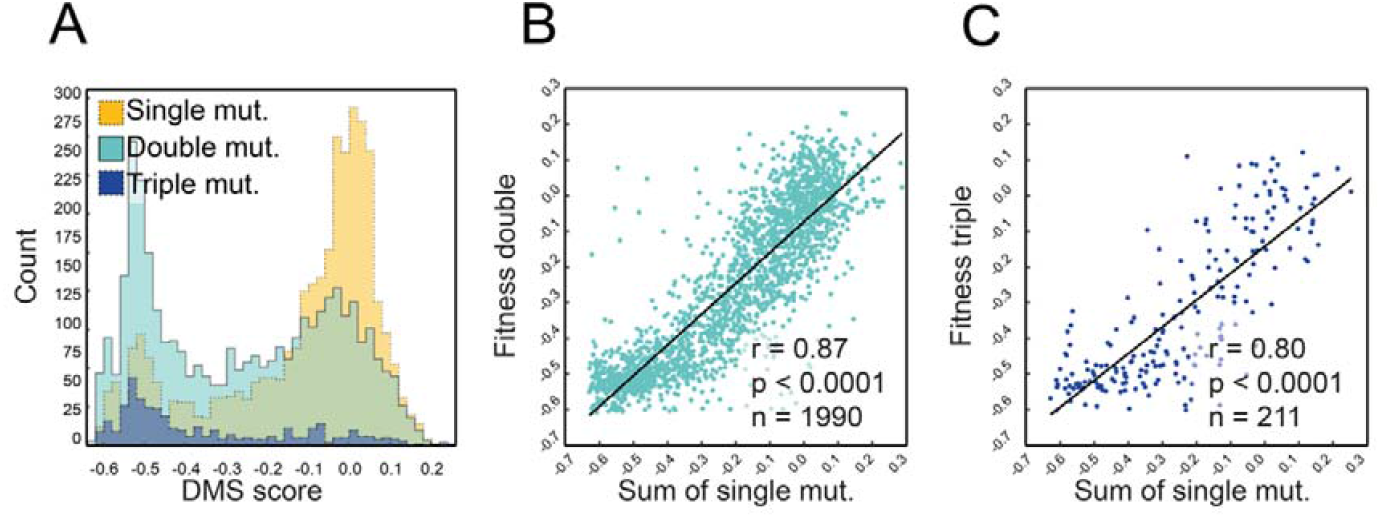
Combinatorial mutant analysis. **A)** Histograms of DMS fitness scores, grouped by number of mutations. Single mutants are shown in yellow, double mutants in cyan and triple mutants in dark blue. **B)** Scatterplot showing DMS scores of double mutants (for F_1,2_ > -0.63) vs. sum of corresponding single mutant DMS fitness scores (F_1_ + F_2_) (*r* = 0.87, *p* < 0.0001), indicating low epistasis among the set of double mutants. **C)** Scatterplot showing DMS scores of triple mutants vs. sum of corresponding single mutant DMS fitness scores (F_1_ + F_2_ + F_3_) (*r* = 0.8, *p* < 0.0001).

In order to study further the effect of combining mutations, we generated a small library of 15 combinatorial variants derived from the 5 best expressing single mutations in the library (**Table S6**). Along with the R32C disulfide bond mutant, we created different sets of double, triple, quadruple and quintuple mutants, and measured their expression level on the yeast cell surface as monogenic cultures. The featured mutations are listed in **Figure 6A** and mapped on the 3D structure in **Figure 6B**. Endpoint residues shown are not obtained from crystal structures but are simply modeled in Pymol software to provide an overview. Mutant ⍰-carbons are all more than 7 ⍰ away from each other and obtain all a score of over 0.1 in the DMS experiment. As expected, and discussed earlier, all highly stabilizing mutations lie on the protein surface, hence not interfering with the hydrophobic core. Monogenic fitness scores for the combinatorial library members were measured and are presented in **Figure 6C**, alongside the single R32C mutation. The color code from **Figure 6A** is used to indicate which mutations are contained in the combinatorial mutants presented in **Figure 6C** (color heights are not quantitative).

**Figure 6.**
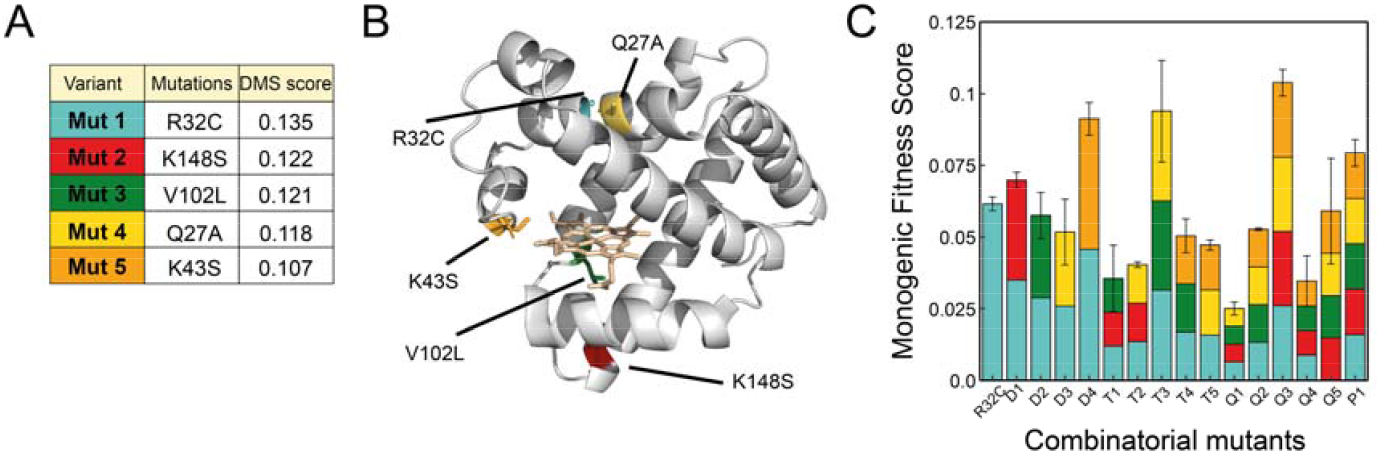
**A)** Table of highly expressing mutants selected for combinatorial analysis. **B)** 3D structure with selected mutations (PDB: 3RGK). Endpoint residues are displayed as sticks and color-coded according to table in A) **C)** Relative fitness of combinatorial mutants measured in monogenic culture. Different colors symbolize which mutations are contained within the combinatorial variants (sizes of colored sub-bars are not quantitative).

All variants show a higher expression level and hence higher thermostability than WT. Several of the combinatorial variants that we tested exhibited higher stability than the R32C disulfide bond reported above. Amongst the double mutants D1-D4, only D4 (K43S & R32C) presents a significantly higher expression level. The triple and quadruple mutants are generally less well expressed than R32C alone, with the exceptions of T3 and Q3, both of which featured a Q27A substitution. We found that the combination of L102V (green) and K148S (red) were detrimental for stability, with the three lowest expressed variants containing those mutations. Taking into account the 3D structure in **Figure 6B**, one could assume that the close 3D distance between the two exchanged residues in neighboring helices might negatively impact the otherwise stabilizing effects of the mutations individually.

Variant P1 contains a combination of all the chosen mutations, and provides an example of how additivity no longer holds for higher order mutants. The variant Q5 contains all the chosen mutations except the R32C disulfide bond. Interestingly we observed an expression level similar to the R32C mutation alone. This again shows the massive impact on stability of this mutant alone, which was further visualized in a correlation matrix uncovering the individual contributions of each mutation (**Figure S5**). We note that that these mutations have not been tested for their compatibility with the physiological function of molecular oxygen binding.

Next, we investigated the effect of multiple mutations on thermostability of soluble proteins. We purified the best 3 (D4, T3 and Q3) variants as determined by the on-yeast expression measurements, and measured their denaturation temperatures via NanoDSF (**Figure S6**). Based on non-reducing SDS page gels, we observed successful disulfide bond formation and high purity for all variants. We found increased stability with measured T_m_ values of 84.85, 89.18 and 84.47 °C for D4, T3 and Q3, respectively. Therefore, in accordance with the yeast surface display expression level measurements, we found higher thermostability for all three combinatorial mutants. However, while for variants D4 and Q3, we detected an increase of 13.51 and 13.13 °C over wild type, respectively, we could measure a massive increase of T_m_ for variant T3, with a melting temperature of more than 17 °C higher than WT.

Compared to the single R32C variant, variants D4 and Q3 are hence only marginally improved. Other recent work from the protein design field presented stability enhancing mutations for hMb using ProteinMPNN^38^. Interestingly, in that work the most stable variant described also features mutation Q27A, along with a mutation of position V102 to an isoleucine, which is closely related to the leucine introduced in our variant T3^39^. However, an important difference is that the ProteinMPNN neural network predicted sequences with ∼41-55% sequence identity while in our case only 3 positions in hMb have been mutated. The variants reported here therefore share much higher sequence identity with parental hMb (98% similarity) while achieving comparable stability enhancements of more than 15 °C.

## Discussion

In this report, we used comprehensive DMS by yeast surface display to identify stabilizing mutations in hMb. We demonstrate correlation of yeast display expression level with thermostability of hMb variants when expressed and tested as soluble proteins. This finding was further supported through predictive computational folding stability models like FoldX and ThermoMPNN. Our comprehensive DMS-derived fitness landscape of single point mutations of hMb showed structural trends and tolerance for mutations in specific regions spread across the entire sequence length, allowing for discovery of crucial sites for stability. We demonstrated how alpha helix pitch could be measured from mutational stability data, and further discovered a highly stabilized interhelical *de novo* disulfide bonded variant of hMb.

We studied the effects of combining single mutations and showed that mutations contributed to expression and thermostability stability in a mostly additive manner for small numbers of mutations, however these linear trends deviated for higher variants containing more than three mutations. Making use of this knowledge, we were able to design a new-to-nature hMb variant, with significantly improved thermostability (T_m_ = 89.18°C), representing >17 °C improvement in T_m_ from wild type. Together, these findings highlight the power of DMS in understanding protein stability and accelerating protein engineering campaigns with broad applications in industry and biomedicine.

## Author contributions

CK: Conceptualization, methodology, investigation, formal analysis, writing (original draft); OP: Methodology, investigation, and data analysis; RV: Conceptualization, methodology, writing (review & editing); MAN: Conceptualization, methodology, writing (review & editing), funding acquisition.

## Supplementary Materials

Supplementary materials are available for this article and include detailed materials and methods, Figures S1-S6, and Table S6.

## Acknowledgements

This work was supported by the University of Basel, ETH Zurich and the Swiss National Science Foundation (200021_191962). The authors further thank Vanni Doffini, Sean Boult and Jaime Fernandez de Santaella for helpful discussions.

## Data availability

The data necessary to reproduce the presented results, including raw sequencing files, scripts, custom computer codes as well as other raw data associated with figures, are available at: https://zenodo.org/records/10658344. Additionally, an interactive colorblind accessible heatmap is provided in the repository.

