## Supplementary Information for "Deep Mutational Scanning Reveals a De Novo Disulfide Bond and Combinatorial Mutations for Engineering Thermostable Myoglobin"

### Materials & Methods

#### Materials

Restriction enzymes, DNA ligase kit and calf intestinal phosphatase (CIP) were purchased from New England Biolabs (Beverly, MA). GelRed^TM^ dye nucleic acid gel stain was purchased from Biotium. Gene Morph II random mutagenesis kit (epPCR) was obtained from Agilent Technologies (Santa Clara, CA). The primary and secondary antibodies were purchased from Thermo Fisher Scientific (Waltham, MA). Custom oligonucleotides were obtained from Microsynth (Balgach, CH). The DNA miniprep, gel purification and PCR purification kits were purchased from Thermo Fisher Scientific. Chemicals, if not indicated otherwise, are from Sigma Aldrich (St. Louis, MO). Sodium alginate was purchased from Duchefa Biochemie (Haarlem, Netherlands).

#### Myoglobin gene cloning

The homo sapiens myoglobin (Mb) gene was obtained from TWIST Bioscience codon optimized for *Saccharomyces Cerevisiae*. The gene was cleaved in a double digest with restriction enzymes EcoRI and XhoI and cloned in a pYD1 expression vector. Sequence was confirmed by sanger sequencing and the plasmid featuring the myoglobin gene transformed in yeast strain EBY100 by a lithium acetate transformation protocol (40). Positive colonies were selected on synthetic defined (SD) agar plates lacking tryptophan (-TRP) supplemented with glucose 2% (wt/vol) and ampicillin.

#### Construction of barcoded myoglobin mutant library

For mutagenesis of the human myoglobin wild type gene, we used a plasmid based one-pot mutagenesis protocol which was previously described (17). Since higher efficiency was observed for smaller plasmids, we transferred the expression cassette from the pYD1 (5446 bp) to a smaller puc19 plasmid (3647 bp). The puc19 plasmid contains two BbvCI restriction sites up- and downstream of the expression cassette. The original protocol was slightly modified by adding an extra step after the first incubation with Nt.BbvCI enzyme and exonuclease. The reaction mixture was followed by incubation with 10 units of quick CIP phosphatase (New England Biolabs) at 37°C for 20 min, with subsequent incubation at 80°C for 20 minutes. With this step, the removal of 5’ phosphate from nicked and partially degraded strands of wild type gene is promoted and hence limits the reformation of closed wild type plasmids in the following amplification and ligation steps. Then, nested NNK codon containing primers were used for synthesis of the mutated strand targeting codon triplets 1 to 154 (full gene) of the myoglobin wild type gene. Primers were designed with the script create_primers.py from (17) whereas the length of the primers was adjusted to reach a similar melting temperature around 60 °C for all primers. Designed primers were ordered from Integrated DNA technologies in a 96 well plate format and mixed in equimolar ratios to the final concentration of 10 µM. With the mixed primers, the site saturation protocol was performed as described in (17). After obtaining the plasmid based mutation library, molecular barcodes were introduced through PCR and hence linked to each of the amplified Myoglobin variants. PCR was accomplished by using primers P1 and P2 shown in **Table S1**, adding a 15 N barcode in front of the galactose promoter sequence. The amplified and barcoded library was cloned in a pYDKan backbone, by HiFi DNA Assembly (New England Biolabs) and using the vector plasmid which was linearized by BbVCI/PmeI double digestion. After column purification (Zymo-Spin I, Zymoresearch), the HiFI assembly product was then transformed into electrocompetent XL1-Blue (Agilent) cells by electroporation (Gene Pulser, Biorad).

Successfully transformed cells were selected for in LB agar plates supplemented with (50 μg/ml) Kanamycin and the size of the library was estimated by counting colony forming units on plates with serially diluted fractions of the library. The total size of the barcoded library was estimated to be 300,000 variants. After incubation at 37°C for 20 hours, the colonies were scraped from the plates and resuspended in liquid LB Kan, further grown for 2 hours and then plasmids extracted by standard Miniprep method. 10 μg from the plasmid library pool was introduced into the *Saccharomyces cerevisiae* strain EBY100, using the lithium acetate transformation protocol(40). To assess the efficiency of the transformation, successive dilutions of the transformed material were spread onto agar plates containing -Trp and 2% (wt/vol) glucose. Counting of the resulting colony forming units revealed a transformation efficiency around 2 million colonies. Following the transformation, the yeast mutant library was cultivated for 24 hours at a temperature of 30°C in a liquid medium lacking tryptophan (-Trp), supplemented with 2% (wt/vol) glucose. Subsequently, the culture was diluted to an OD_600_ of 1 and transferred to fresh -Trp medium with 2% (wt/vol) glucose to minimize the chance of obtaining multiple transformants. This culture was then allowed to grow for an additional 16 hours at 30°C. Finally, portions containing approximately 5 *10^7 cells were prepared and suspended in a solution of 25% (vol/vol) glycerol before being preserved at -80°C. Plasmid maps are provided at: DOI: 10.5281/zenodo.10658344.

**Table S1)** Primers used for linkage of molecular barcodes to the myoglobin mutant variants. NNK nested primers used for the generation of the library are provided at: DOI: 10.5281/zenodo.10658344

| Barcoding Primer P1 | CGATTTTGTTACATCTACACTGTTGTTATCAGATCAGCGGGTTTAAAC |
| --- | --- |
| Barcoding Primer P2 | GCGCGGCCTTTTGCCCTGCAGGCCNNNNNNNNNNNNNNNAGGGAACAAAAGCTGG CTAGTACGG |
| PacBio 1 | TTACGGTTCCTGGCGGCCGCG |
| PacBio 2 | GTCGATTTTGTTACATCTACACTGTTGTTATCAGATCAGCGGG |
| Illumina_for1 | TCGTCGGCAGCGTCAGATGTGTATAAGAGACAGGGCCTTTTGCCCTGCAGGC |
| Illumina_for2 | TCGTCGGCAGCGTCAGATGTGTATAAGAGACAGTGGCCTTTTGCCCTGCAGGC |
| Illumina_for3 | TCGTCGGCAGCGTCAGATGTGTATAAGAGACAGATGGCCTTTTGCCCTGCAGGC |
| Illumina_for4 | TCGTCGGCAGCGTCAGATGTGTATAAGAGACAGCATGGCCTTTTGCCCTGCAGGC |
| Illumina_rev1 | GTCTCGTGGGCTCGGAGATGTGTATAAGAGACAGGGAGGAGAGTCTTCCTTCGGAGGG |
| Illumina_rev2 | GTCTCGTGGGCTCGGAGATGTGTATAAGAGACAGTGGAGGAGAGTCTTCCTTCGGAGGG |
| Illumina_rev3 | GTCTCGTGGGCTCGGAGATGTGTATAAGAGACAGATGGAGGAGAGTCTTCCTTCGGAGGG |
| Illumina_rev4 | GTCTCGTGGGCTCGGAGATGTGTATAAGAGACAGGATGGAGGAGAGTCTTCCTTCGGAGGG |

In order to limit the size of the library and increase the confidence in barcode assignment from the PacBio-generated lookup table, we capped/bottlenecked the library to 100,000 cells. This was done on a FACS Sony SH800S Sorter by simply gating for viable single cells and sorting 100,000 events in growth media. This culture was grown for 20 h overnight and stored as glycerol stocks at -80°C.

#### PacBio long read sequencing

In order to link the molecular barcodes to their respective hMb variant sequence, Pacbio long read sequencing was used. The capped library glycerol stock was thawed and used to inoculate 40 mL -TRP and 2% (wt/vol) glucose at an initial OD of 0.1 and allowed to grow for 24 hours. OD then was around 6.8 and 1 mL (4 samples) of the cultures were collected, spun down, and resuspended in 250 µl of Miniprep Resuspension buffer 1 (GeneJET, ThermoFisher). Then 4 µl of Zymolyase was added (Zymoresearch) and incubated for 2 h at 37°C with shaking at 1,000 rpm. Then the standard protocol as suggested in the kit was followed, whereas the final elution volume was 10 µl of prewarmed nuclease free sterile water. 9 µl of the eluted DNA were used as template for a standard PCR using the Q5^®^ High-Fidelity 2X Master Mix (New England Biolabs) and 26 cycles of amplification. In total, 4 PCR reactions were conducted from 4 individual zymopreps and all amplicons were loaded on an agarose gel. Purified bands were digested in a double digest with restriction enzymes Pme1 and Not1. Then, amplicons were purified and concentrated with a DNA clean and concentrate kit (Zymo research). The purified DNA was then used for SMRT bell ligation and run on a Pacific Biosciences Sequel IIa sequencer with a movie time of 15 hours. Resulting sequences were filtered for quality score Phred 20 and higher. Additionally, sequences were filtered for a maximal length of 1800 bp (amplicon length is 1569). Then, the computational workflow already established and reported in (5) was used to extract the barcode and assign it to a respective mutation. We used Minimap2 (18) to map the long read sequences to a reference file with the wild type sequence of the expression vector PCR amplicon. Then, a C script was used to process the same file output of the mapped sequences using a custom library file (libscodon.h). The script extracts the 15 base pair molecular barcode and its quality along with the mutational sequence for every read. The output of this step is a first version of the lookup table. Errors such as barcodes assigned to multiple mutants in this raw version of the LUT are then purged in a python script collapsing all the reads, and filtering them for their confidence. Barcodes with the highest number of reads linking the barcode to the mutation are kept and if two barcodes were read the same number of times, the one with the higher quality was kept in the LUT. Then, the LUT was filtered eliminating barcodes that have the wrong length and barcodes that were read less than twice.

**Table S2)** Computational workflow. Steps and respective commands to map the PacBio reads to the reference and extract the molecular barcode to generate the final look up table (LUT). The raw sequencing files as well as the computational scripts are provided at: DOI: 10.5281/zenodo.10658344.

| **Step** | **Tool** | **Command** |
| --- | --- | --- |
| Mapping of the PacBio read to the reference file. | Minimap2  (version 2.19) | **minimap2 --cs -ax map-hifi** Myo_ref.fa pacbio.fastq **>** aln.sam |
| Extracting barcode sequence and quality along with resp mutations and indels. | ppba C script  libscodon.h library | **./ppba** Myo_ref.fa aln.sam **>** pre_lut.tsv |
| Filtering and cleaning of LUT | generate LUT.ipynb  Python script |  |

#### Expression and labeling of myoglobin barcoded library

Aliquoted glycerol stocks of the library were thawed on ice and resuspended in liquid -TRP medium supplemented with 2% (wt/vol) glucose and Ampicillin 100 μg/ml at a starting OD of 0.1 in 50 mL. After preculture growth for 30 hours with shaking (180 rpm), appropriate volume of cells were spun down and resuspended in 100 mL at an OD 0.4 in -TRP liquid induction medium with 1.8% (wt/vol) galactose, 0.2 % glucose and Ampicillin, buffered with 100 mM sodium phosphate buffer pH 7. Cells were grown at 20°C for 48 hours with shaking at 180 rpm. Then, the culture was spun down, washed once, and resuspended in PBS BSA 0.1% (wt/vol). For cell staining, 4 *10^6^ cells were collected in a round bottom 96 well plate and incubated in 100 ul of 1:500 dilution of primary Anti Histag Antibody (stock concentration of 1.2 mg mL^−1^). Afterwards, cells were washed again with 200 ul of PBS BSA 0.1% (wt/vol) and incubated for 20 min in ice cold dilution (1:500, stock concentration 2 mg mL^−1^) of secondary goat anti-mouse antibody conjugated with Alexa Fluor 488. Then, cells were washed again twice with cold PBS BSA 0.1%.

#### Fluorescent activated cell sorting for deep mutational scanning

The capped, expressed and stained library was sorted on a BD FACS Melody. After labeling for expression, the library was loaded on the Cell sorter and gates were applied along the Alexa fluorophore 488 (BL1 channel) axis capturing the whole negative population (∼ 40%) in gate 1(-) and dividing the remaining population in three equi percentage gates 2(+), 3(++) and 4 (+++) . The sorting experiment was repeated twice independently on the same day to achieve biological replicates, sorting each time at least 2.5 Million cells covering the library 25 times. Additionally, we sorted 1000 cells from a separately grown expression culture of wild type with a known barcode in every sort tube. This internal control allows for monitoring any bias in growth conditions in between tubes.

The gated populations were sorted in -TRP media supplemented with 1 % BSA to facilitate pelleting of the cells. Cells were spun down and resuspended in 10 mL -TRP medium with 2% glucose and ampicillin. The cultures were let recover and grow for 48 hours at 30°C with shaking at 180 rpm. Then, aliquots of 5 *10^7 cells were stored as glycerol stocks at -80°C.

#### Illumina barcode sequencing

In order to prepare the sorted bins for high throughput sequencing, glycerol stocks (2 per bin) from both biological replicates were thawn for 5 minutes on ice and spun down at 13000 g for 1 min in a benchtop centrifuge. Then, the supernatant was removed and 250 ul of resuspension solution from bacterial miniprep kit (GeneJET Plasmid Miniprep Kit, Thermo Fisher) were added for resuspension of the pellet. Per tube, 4 ul of Zymolyase was added, and let incubate for two hours at 37 °C with shaking at 1000 rpm. After visual inspection for successful lysis of yeast cell walls (solution appears clear), 250 ul of lysis buffer and subsequently 350 ul of neutralization buffer was added, following the standard protocol for Plasmid miniprep. For better coverage of binned barcodes, two Zymoprep tubes per bin were combined in one miniprep column. After conducting the washing steps, final elution was done in 15 ul of nuclease free water. Those 15 ul then served as template for the PCR reaction, amplifying the 15 N barcode containing region in a standard 50 ul PCR reaction (NEBNext Ultra II Q5 Master Mix, New England Biolabs). 25 ul of Master Mix were added to 15 ul of template DNA with 5 ul of each primer (final conc 1 uM). For forward and reverse primers used to amplify individual binned genetic information, we used staggered primers (Ilumina_for/rev 1-4, **Table S1**), in order to provide different starting nucleotides during illumina sequencing in an else very similar amplicon. Primers were designed for use with the Illumina Nextera indexing library prep.

The amplicons for all bins were then loaded on a standard 1 % (wt/vol) agarose gel to verify correct size and then purified form cut gel using a GeneJET Gel Extraction Kit (Thermo Fisher). After cleaning and concentrating the DNA further using a DNA Clean and Concentrator-5 kit (Zymoresearch), we determined DNA concentration. In a second PCR (**Table S4**) using 1 ng of template DNA, the Nextera indexing sequence was added for all of the bins. Samples were purified with AMPure XP beads, pooled and sequenced through Novaseq 6000 Illumina sequencing.

**Table S3)** PCR conditions for barcode amplification

| **Step** | **Temp. [°C]** | **Time [s]** | **Cycles** |
| --- | --- | --- | --- |
| Initial Denaturation | 98 | 30 | 1 |
| Denaturation | 98 | 10 | 20 |
| Annealing | 72 | 30 |  |
| Extension | 72 | 10 |  |
| Final extension | 72 | 120 | 1 |

**Table S4)** PCR conditions for Illumina Nextera Indexing

| **Step** | **Temp. [°C]** | **Time [s]** | **Cycles** |
| --- | --- | --- | --- |
| Initial Denaturation | 98 | 30 | 1 |
| Denaturation | 98 | 30 | 8 |
| Annealing | 55 | 30 |  |
| Extension | 72 | 30 |  |
| Final extension | 72 | 120 | 1 |

A computational pipeline was established to process the obtained Illumina reads in order to receive the information about abundance of corresponding mutations in each bin. By using the BBMap algorithm and providing the illumina reference file containing a 15 N barcode, the sequences were mapped and alignment information stored in a sam file (19). Barcodes were then extracted with the pib.c script and linked through their corresponding mutation by running the rib.c script which searches for the barcodes and assigns tags according to the description in **Table S5**. Finally, all barcodes present in the LUT were sorted and grouped by identity, revealing the total number of each variant in each bin.

**Table S5)** Pipeline for Illumina read processing. All bins were processed separately through the individual owing steps. The raw sequencing files as well as the computational scripts are provided at: DOI: 10.5281/zenodo.10658344.

| **Step** | **Tool** | **Command** |
| --- | --- | --- |
| Sequence alignment of illumina reads to reference file. | BBMap  cite | bbmap.sh in=*.fq ref=reference.fa out=*.sam |
| Extracting barcode sequence and respective quality score for the illumina reads | pib.c script  (process illumina barcode) | pib *.sam > *.fq |
| Align the extracted barcodes to the previously processed look up table and assign tags (0 = not found, 1 = found, 2 = Read quality below Q20, 3 = Barcode length deviates from 15). | rib.c script  (read illumina barcode) | rib -t lut.ts -q 20 *.fq > *.tsv |
| Filtering for found variants, sort them alphabetically and group them by identity counting the number of barcodes per bin. |  | grep ^1 *.tsv\|sort\|uniq -c\|sed -E 's/^ *//; s/ /\t/' > t1sct_*.tsv |

In a jupyter notebook (ill_tag1_bins) , the grouped barcodes from all the bins were merged into one file, which was then used as the input file of the further analysis in the python script.

#### Fitness score quantification

The files containing the abundance info for all barcodes in all bins was loaded in a jupyter notebook, processing the data for the generation of a fitness score for all mutants. Raw read numbers were transformed into numbers of sorted cells. This was done by multiplying the variant read count by the amplification factor $(\frac{1000}{Rwt}$) that could be estimated by the read count of the known WT barcode variant $Rwt$, which was spiked in a thousand times (1000 cells) in every post-sort recovery tube.

$Cv =Rv(\frac{1000}{Rwt}$)	(1)

,whereas $Cv$ is the cell count and $Rv$ the Illumina read count for a variant in a bin and $Rwt$ the Ilumina count of the internal control barcode.

Then, for every barcode, a fitness value was determined by calculating the expected value of the fluorescent intensity across all gated bins. The total cell number for all barcodes was multiplied by the weights given through the MFI in every bin, and hence a weighted mean was determined.

$\beta= \frac{\sum_{i=1}^{gates} \omega_{i}\times\%c_{i}}{\sum_{i=1}^{gates} \%c_{i}}$	(2)

Finally, the stability fitness score was defined as the log2 of the ratio of weighted mean per variant over weighted mean of the wild type.

$F_{Exp} = log2(\frac{\beta_{Var}}{\beta_{WT}}$)	(3)

For the biochemical analysis of the library (**Figure 2**), scores from barcodes coding for the same mutation were averaged and we only considered mutations that were covered by at least 15 cells among all bins.

#### Monogenic validation of fitness scores

In order to determine the fitness score for monogenetic cultures of the variants, yeast competent cells were transformed with the respective plasmids and variants expressed on the surface as described above. Then, cells were stained with the same antibodies and protocol as used in the DMS experiment and loaded on the Attune Flow cytometer. Again, 4 gates were applied capturing all singlet cells. The gates were set on the wild type population to closely match the percentages obtained for wild type sequences in the DMS experiment. Then, the gates were maintained for all variants studied and percentages as well as median fluorescent intensities per gates extracted in independent triplicates. For each variant, the percentage of cells falling in each gate was multiplied with the mean MFI for the same gate over all variants. We determined a weighted mean for every variant by using the percentage of cells falling in each gate as weight, similarly to the procedure used in generating the high-throughput DMS score (Equation 2, where MFI corresponds to the mean of the per gate overall variants).

To set the calculated score in relation with the wild type, we report the final score as the log2 values of the ratio (Equation 3).

#### Soluble expression and purification of myoglobin variants in *E. Coli*

Genes coding for the selected variants were cloned in pet28 bacterial expression vector and after Sanger sequencing confirmation transformed in BL21(DE3) or Origami2 competent cells and selected on respective LB plates. A single colony was used to inoculate 3 mL of LB Kan and grown overnight at 37°C. (200 rpm) Then the pre-culture was diluted 1:100 in a total volume of 50 mL and let grow until the culture reached OD 0.5. Then, 0.5 mM of IPTG was added to induce expression and the culture was grown at 25°C for 20 hours with shaking (200 rpm). Then to harvest the now reddish cells, they were spun down at 4000 rpm for 10 min and then resuspended in 10 mL lysis buffer (50 mM TRIS, pH 8.0, 50 mM NaCl, 0.1% (v/v) Triton X-100, 5 mM MgCl2). Then, cells were lysed by sonication for 10 minutes (Sonifier cell disruptor Branson Digital Sonifier, USA) and spun down at 18 000 g for 30 min at 4 °C. The supernatant was loaded on a gravitational flow column packed with 0.5 mL of HisPur Ni-NTA resin. The loaded column was washed with 6 column volumes of PBS with 50 mM imidazole and eluted in 2 mL 500 mM Imidazole PBS. Elution fraction was concentrated in vivaspin concentration columns (5 kDa cutoff) and salt was removed by Zeba Spin desalting column (ThermoFisher).

#### NanoDSF procedure

For estimation of thermostability of the purified soluble myoglobin variants, the proteins were diluted in PBS to a final concentration of 0.5 mg/ml and loaded on a Prometheus NT.48 NanoDSF (NanoTemper Technologies, Munich, Germany) in standard capillaries unsealed (Cat# PR-C002). Samples were melted at 2 °C/min and intrinsic fluorescence detected from 20 to 95 °C at a 100% excitation intensity. Melting temperatures T_m_ were defined as the peak of the first derivatives of the ratio (330 nm/350 nm) from three technical replicates.

#### FoldX studies

For the computational analysis of the predicted ΔΔG values, we used the foldX stability algorithm (32)**.** First, the PDB file for human myoglobin including heme cofactor (3RGK) was processed with the repair function of the FoldX suite (RepairPDB). This is strongly suggested by the authors of FoldX and identifies energetic clashes and finds rotamer combinations to yield energy minima. The PositionScan command then mutates all amino acids to all other natural amino acids and calculates a predicted value for the change of energy in between native and unfolded confirmation for wild type and all introduced mutations. The change of the change of Gibbs free energy (ΔΔG) is then calculated according to:

ΔΔG = ΔG_Mut_ - ΔG_WT_

The ΔΔG scores were then averaged overall mutations at one position and compared to the experimental expression level fitness scores. The raw sequencing files as well as the computational scripts are provided at: DOI: 10.5281/zenodo.10658344.

#### ThermoMPNN ΔΔG prediction

For an alternative stability change prediction of introduced point mutations, we turned to ThermoMPNN, a deep neural network which was recently released (33). We used their Colab implementation of the neural network and imported the human Mb structure (PDB: 3RGK, n = 149).

We then exported the output data and merged it with the DMS stability scores. The ΔΔG scores were again analyzed individually or averaged overall mutations at one position and compared to the experimental expression level fitness scores. The raw sequencing files as well as the computational scripts are provided at: DOI: 10.5281/zenodo.10658344.

#### Mass spectrometry validation of disulfide bond

Myoglobin variant R32C was purified along with WT from oxidizing strain Origami2 as described above. Then, buffer was exchanged to water and samples were loaded on a LC-MS (LC: Thermo FIsher Ultimate 3000 UPLC, MS: Bruker maxis 4G ESI-Q-TOF) at 0.2 mg/ml supplemented with 0.1 % (v/v) of formic acid.

### Supplementary Figures

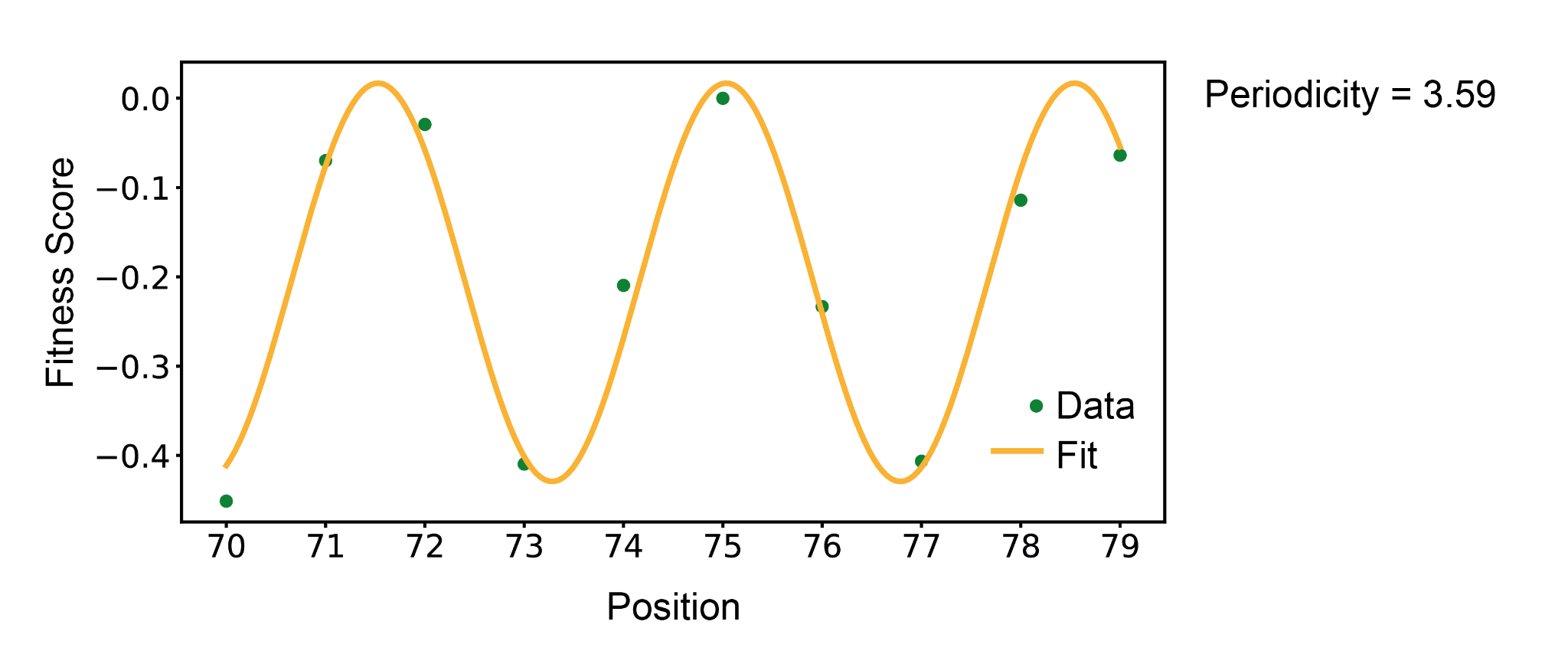

**Figure S1.** Averaged fitness scores for positions 70-79 and fitted periodic function with a periodicity = 3.59, matching the pitch of an alpha helix.

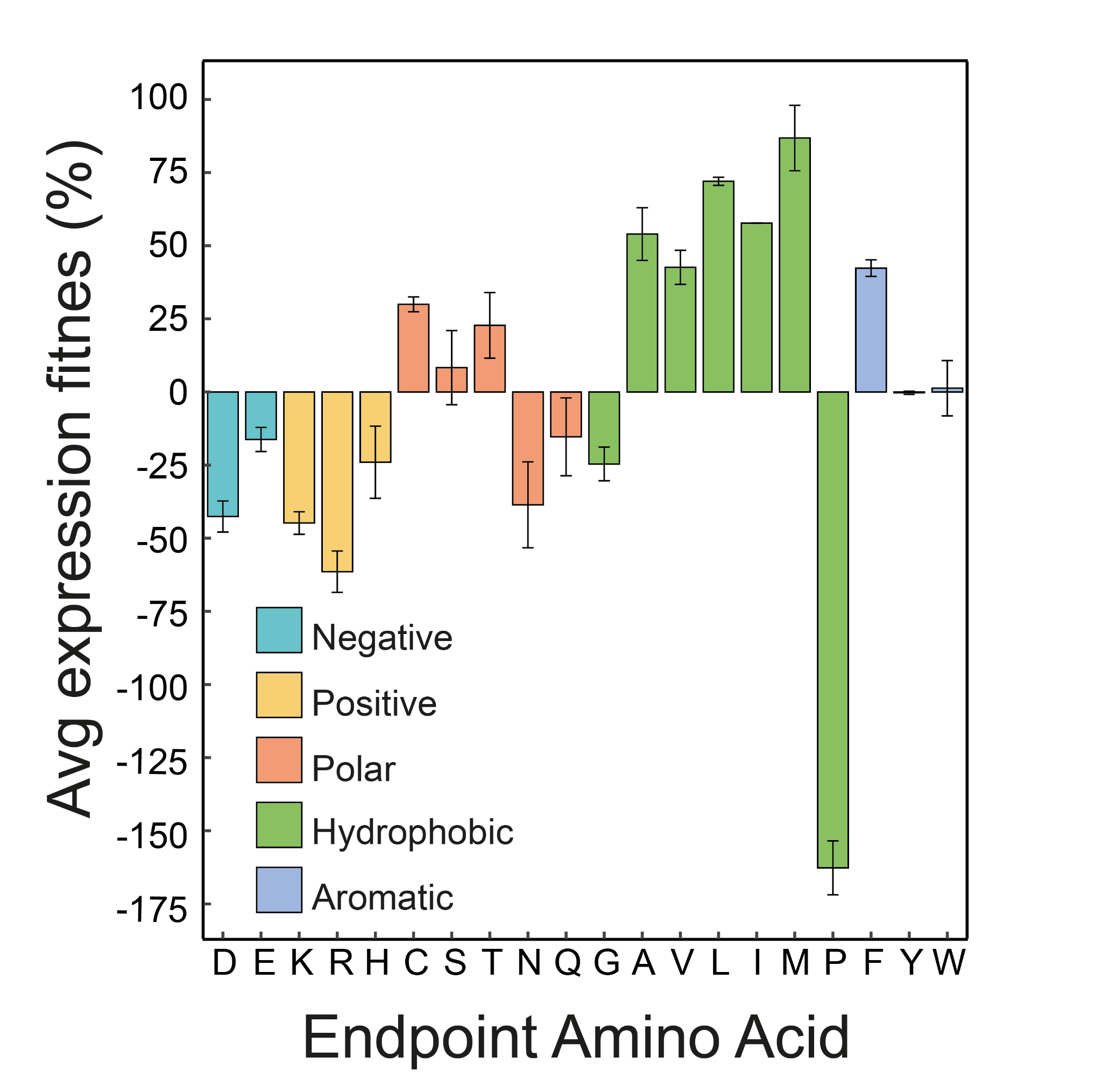

**Figure S2.** Averaged fitness impact of grouped endpoint amino acids, expressed as percent change difference against overall endpoint fitness.

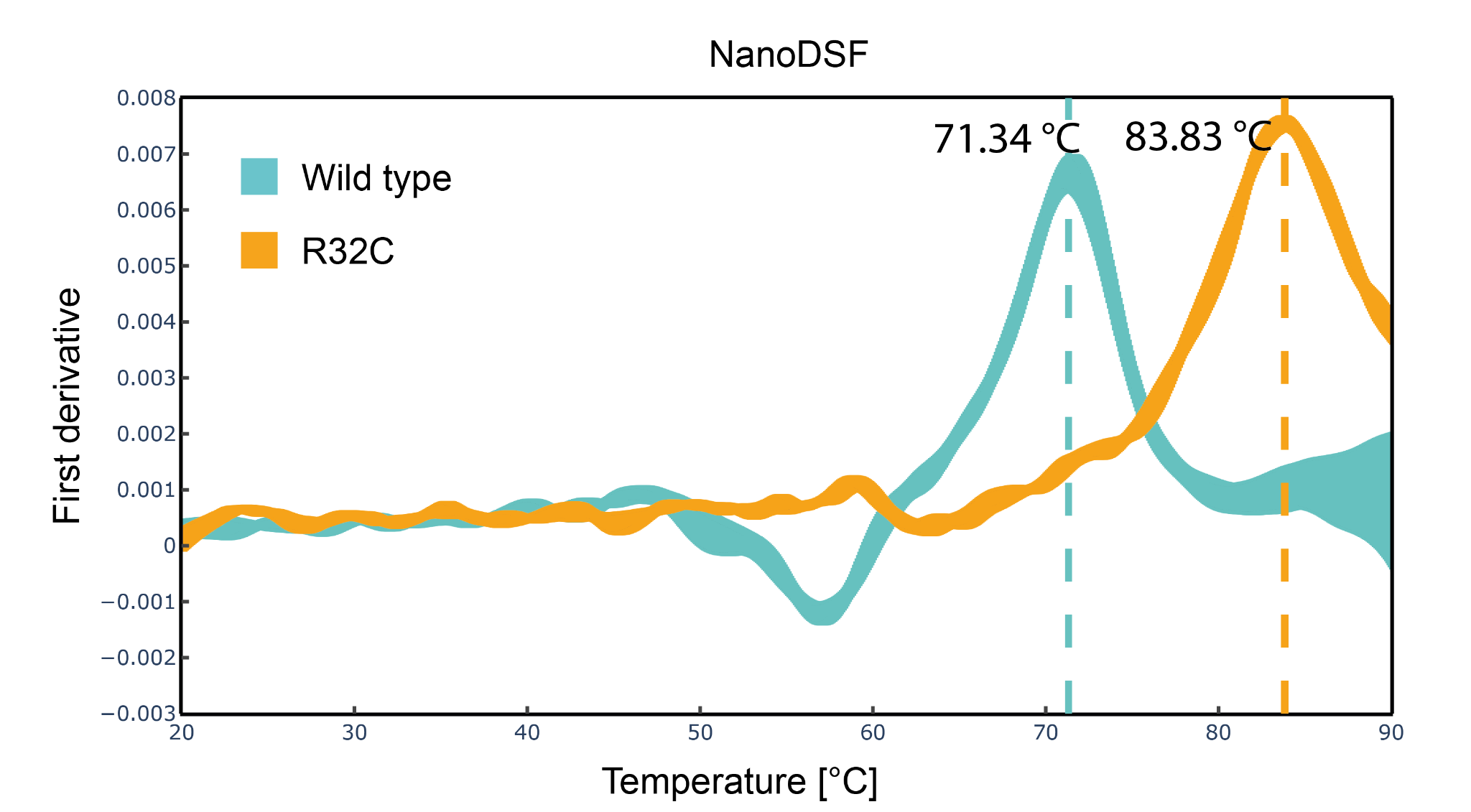

**Figure S3.** First derivatives of the melting curves for Myoglobin wild type sequence and R32C variant by NanoDSF, averaged over technical triplicates.

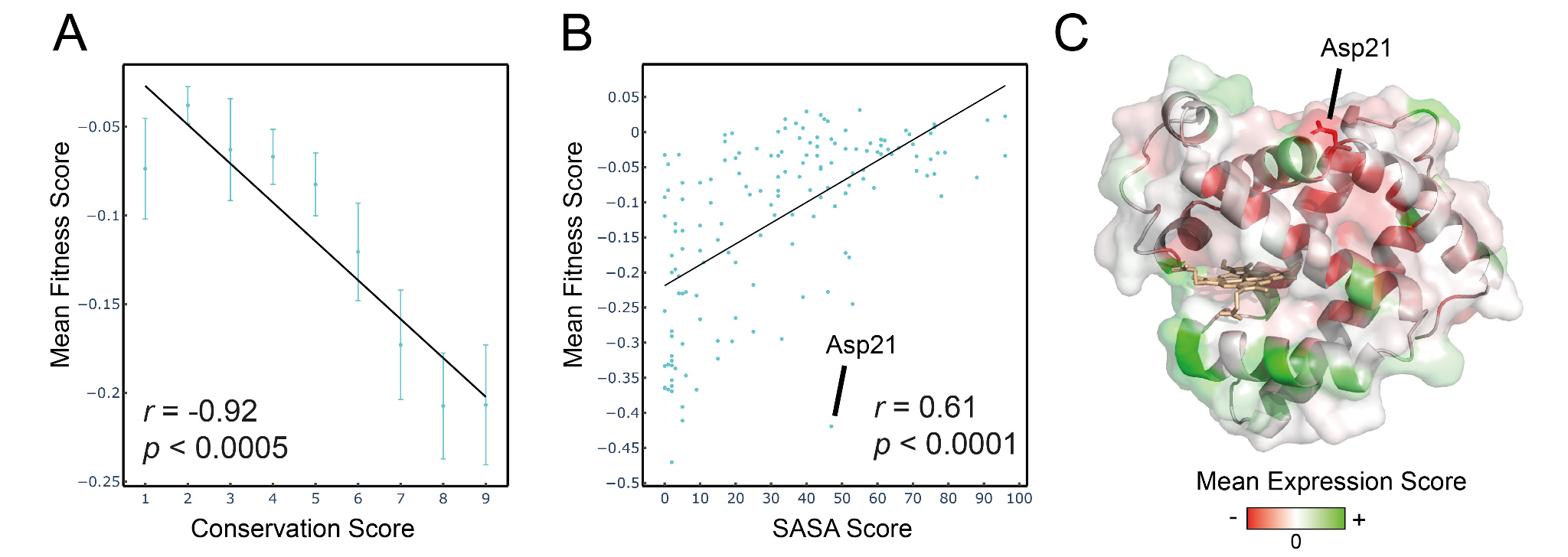

**Figure S4.** A) Mean fitness per position grouped by conservation score. Error bars are standard errors of all positional fitness scores per conservation score unit. B) Correlation of mean fitness and solvent accessible surface area (SASA) score per position. C) Myoglobin 3D structure (AF-P02144-F1), colored by mean per position fitness. The low tolerance residue Asp21 is highlighted in stick representation. The raw data files are provided at: DOI: 10.5281/zenodo.10658344.

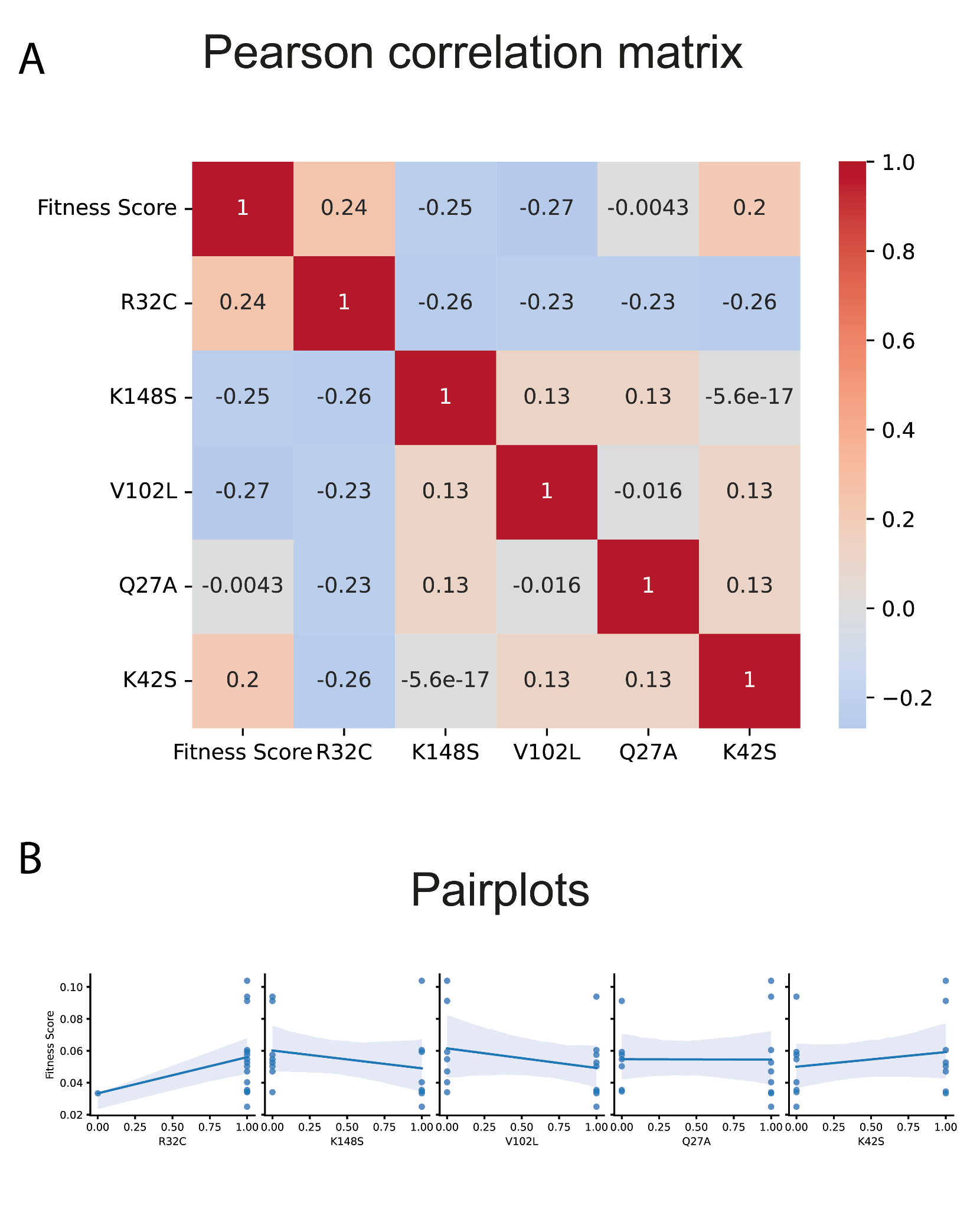

**Figure S5.** Contribution of individual mutations to fitness scores visualized as Pearson correlation matrix (A) or pairplots (B).

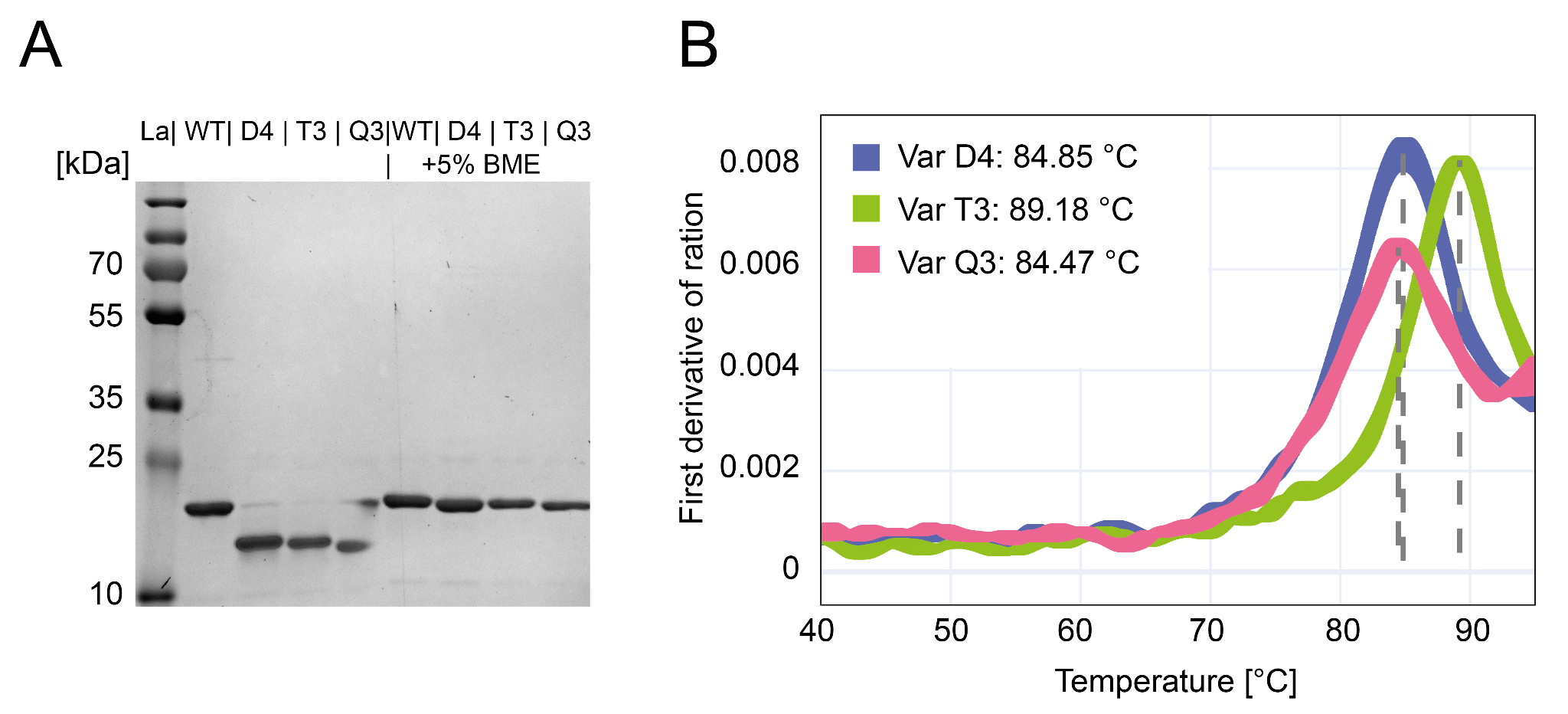

**Figure S6. A)** SDS page of purified high stability variants along with WT control. The apparent smaller size is a consequence of the disulfide bond and is not observable when run under reducing conditions. Leaking of β-mercaptoethanol (BME) to neighboring lanes leads to partial reduction in lane of unreduced Q3. **B)** First derivatives of the melting curves for high stability variants by NanoDSF, averaged over technical triplicates.

**Table S6.** Selected mutants for generation of combinatorial library of stabilizing residues.

|  | **Mutation 1** | **Mutation 2** | **Mutation 3** | **Mutation 4** | **Mutation 5** |
| --- | --- | --- | --- | --- | --- |
| **Variant** | **R32C** | **K148S** | **V102L** | **Q27A** | **K42S** |
| D1 |  |  |  |  |  |
| D2 |  |  |  |  |  |
| D3 |  |  |  |  |  |
| D4 |  |  |  |  |  |
| T1 |  |  |  |  |  |
| T2 |  |  |  |  |  |
| T3 |  |  |  |  |  |
| T4 |  |  |  |  |  |
| T5 |  |  |  |  |  |
| Q1 |  |  |  |  |  |
| Q2 |  |  |  |  |  |
| Q3 |  |  |  |  |  |
| Q4 |  |  |  |  |  |
| Q5 |  |  |  |  |  |
| P1 |  |  |  |  |  |
